# Lateral and longitudinal fish eDNA distribution in dynamic riverine habitats

**DOI:** 10.1101/2020.05.28.120147

**Authors:** Bettina Thalinger, Dominik Kirschner, Yannick Pütz, Christian Moritz, Richard Schwarzenberger, Josef Wanzenböck, Michael Traugott

## Abstract

Assessing the status and distribution of fish populations in rivers is essential for management and conservation efforts in these dynamic habitats. Currently, methods detecting environmental DNA (eDNA) are being established as an alternative and/or complementary approach to the traditional monitoring of fish species. In lotic systems, a sound understanding of hydrological conditions and their influence on the local target detection probability and DNA quantity is key for the interpretation of eDNA-based results. However, the effect of seasonal and diurnal changes in discharge and the comparability of semi-quantitative results between species remain hardly addressed. We conducted a cage experiment with four fish species (three salmonid and one cyprinid species) in a glacier-fed, fish-free river in Tyrol (Austria) during summer, fall, and winter discharge situations (i.e. 25-fold increase from winter to summer). Each season, water samples were obtained on three consecutive days at 13 locations downstream of the cages including lateral sampling every 1-2 m across the wetted width. Fish eDNA was quantified by species-specific endpoint PCR followed by capillary electrophoresis. Close to the cages, lateral eDNA distribution was heterogenous and mirrored cage placement within the stream. In addition to the diluting effect of increased discharge, longitudinal signal changes within the first 20 m were weakest at high discharge. For downstream locations with laterally homogenous eDNA distribution, the signals decreased significantly with increasing distance and discharge. Generally, the eDNA of the larger-bodied salmonid species was less frequently detected, and signal strengths were lower compared to the cyprinid species. This study exemplifies the importance of hydrological conditions for the interpretation of eDNA-based data across seasons. To control for heterogenous eDNA distribution and enable comparisons over time, sampling schemes in lotic habitats need to incorporate hydrological conditions and species traits.

## Introduction

In times of rapid environmental changes there is a growing need for biomonitoring in both terrestrial and aquatic systems (Cardinale et al., 2012; Tickner et al., 2020). Reliable and cost-effective approaches for species detection are thus key for tracking species in time and space and informing conservation and management efforts (Jetz et al., 2019). Molecular methods, like the detection of environmental DNA (eDNA) released by organisms into their environment, have the capacity to accommodate this demand as they are non-invasive, sensitive, and enable the processing of large sample numbers (Barnes & Turner, 2016; Deiner et al., 2017; Thomsen & Willerslev, 2015). In aquatic habitats, eDNA can be used for the monitoring of taxa in both lotic and lentic systems with a focus on endangered or invasive species and flexible year-round application (Beng & Corlett, 2020; e.g. Harper, Griffiths, et al., 2019; Thomsen & Willerslev, 2015).

Rivers and streams contain, absorb, and transport eDNA from aquatic and terrestrial species and are thus ideal for cross-habitat species detection (Deiner, Fronhofer, Mächler, Walser, & Altermatt, 2016; Sales et al., 2020), although the dynamic nature of these ecosystems leads to constantly changing conditions during sampling (Shogren et al., 2017; Willett, McCoy, Taylor Perron, Goren, & Chen, 2014). Nevertheless, aspects such as a species’ upstream distribution limits (Carim et al., 2019; Robinson, de Leaniz, & Consuegra, 2019) or local abundance and its change over time (Doi et al., 2017; Levi et al., 2019; Thalinger, Wolf, Traugott, & Wanzenböck, 2019) have been successfully examined in lotic systems via eDNA. Often, these efforts are combined with traditional monitoring techniques to confirm molecular results and facilitate their interpretation (Evans, Shirey, Wieringa, Mahon, & Lamberti, 2017; Wilcox et al., 2016) and there is generally a good consensus between molecular and non-molecular data.

However, changes in hydrological conditions have profound influence on the distribution and persistence of eDNA in the water column (reviewed by Harrison, Sunday, & Rogers, 2019), which makes the interpretation of local eDNA signals or comparisons between sampling campaigns challenging. The longitudinal eDNA detection probability in rivers depends on dilution, transport, deposition, resuspension, and degradation (reviewed by Harrison et al., 2019). These effects were examined for small streams (Fremier, Strickler, Parzych, Powers, & Goldberg, 2019; Shogren et al., 2017; Wilcox et al., 2016) and large river systems (Deiner & Altermatt, 2014; Pont et al., 2018), revealing the behavior of eDNA as similar to that of fine particulate organic matter. Recently, studies also focused on lateral eDNA distribution, describing a “plume” downstream of the source and gradual lateral homogenization with increasing downstream distance (Laporte et al., 2020; Wood, Erdman, York, Trial, & Kinnison, 2020). Albeit discharge-associated changes in longitudinal eDNA distribution were previously examined for small streams (Jane et al., 2015), changes in the size and shape of eDNA plumes have been neglected so far.

Alpine rivers exemplify the benefits of eDNA-based monitoring as well as the challenges associated with sampling campaigns in such dynamic ecosystems. The prevailing low water temperatures permit only humble population densities and a limited species inventory at risk of biodiversity loss due to increased climate change effects (Settele et al., 2015). Anthropogenic influences in the form of straightened riverbeds, hydropower plants and dams add to this strained situation and intensify the need for monitoring the remaining natural fish populations (Faulks, Gilligan, & Beheregaray, 2011; Fette, Weber, Peter, & Wehrli, 2007). The discharge in these rivers varies with changing seasons from winter drought to high water levels in spring and summer (snow melt and glacial melt), with intermediate conditions in fall. Additionally, the melting processes induce substantial diurnal discharge changes in spring and summer (Bard et al., 2015). Therefore, traditional monitoring via electrofishing is only possible outside of protected periods (e.g. spawning seasons) and at low discharge and turbidity in fall and early winter. eDNA-based approaches can potentially overcome these limitations, but only after accounting for the spatio-temporal dynamics of Alpine rivers and by optimizing sampling schemes and carefully interpreting the obtained results.

In this study, we examined changes in lateral and longitudinal eDNA distribution between seasons in a glacier-fed Alpine river. The experiments were conducted with caged fish and we used species-specific endpoint PCR combined with capillary electrophoresis (celPCR) to investigate how lateral and longitudinal eDNA detection probability and signal strength vary on a small scale (≤20 m) between summer, fall, and winter, (i.e. high, medium, and low water levels) with up to 25-fold variations in discharge. Additionally, the longitudinal change in eDNA signal strength was examined at an intermediate scale (∼1 km) for fish species of different size, but constant total biomass.

## Materials and Methods

### Study site

The three caged-fish trials took place in the Melach, a fish-free, glacier-fed, Alpine river in Lüsens, Tyrol (Latitude 47°7’0,32”, Longitude 11°8’4,92”, WGS1984; Fig. 1) in August (22^nd^ - 26^th^) 2016, November (21^st^ -25^th^) 2016, and September (26^th^ -29^th^) 2017. The Melach shows seasonal and daily fluctuations in discharge and sediment load, typical for glacially influenced rivers in the Alps (Sertić Perić, Jolidon, Uehlinger, & Robinson, 2015). Following the European river zonation, the sampling site is located in the epirhithral of the river at an elevation of 1700 m a.s.l. Permanent fish populations cannot be established in this part of the river due to extreme discharge situations. Additionally, a transverse structure at the downstream end of the examined range prohibited potential upstream migration.

**Figure 1:**
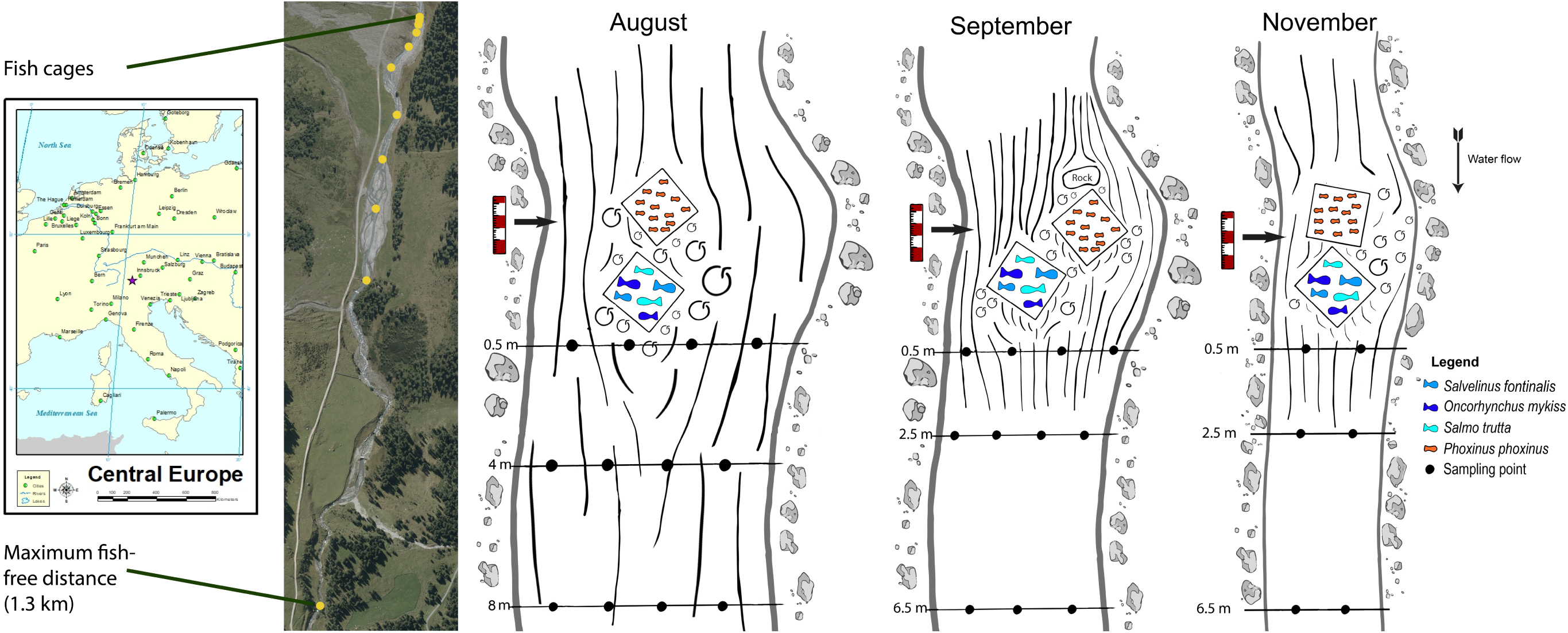
The geographic location of the study region and the aerial view (source: Land Tirol – data.gv.at) of the sampling reach is displayed with yellow dots showing the individual sampling locations at 1.3 km, 555 m, 425 m, 325 m, 225 m, 130 m, 65 m, 33 m, 17 m, 8 m, 4 m, 2.5 m and 0.5 m. Note that downstream distances between 0.5 m and 17 m are not fully resolved in the aerial view. The illustrations show the cage placement in the stream, individual transect samples, and the hydrological conditions prevailing during the August, September, and November trials. Larger flowlines and eddies code for stronger currents. Due to reduced discharge, the relative location and number of eDNA samples were not kept uniform between trials.

### Experimental design

All handling of study animals conformed to Directive 2010/63/EU, and permission for field work was granted by the fishing area manager H. Raffl. Prior to the experiment, the absence of fish was confirmed via electrofishing, starting 1.3 km downstream at the transverse structure. For each of the three trials, two steel cages 1 × 1 × 0.6 m with “mesh” size of 10 mm and 5 mm, respectively, were installed at the same location in full current of the river for the duration of the experimental run (four days), secured against drift, and equipped with a few stones to provide natural structure and potential resting places. Four fish species were used in the experiment: a cyprinid species (Eurasian minnow – *Phoxinus phoxinus* L. 1758) was placed in the cage with the narrow mesh size and three salmonid species (brown trout – *Salmo trutta* L. 1758, brook trout – *Salvelinus fontinalis* M. 1814, and rainbow trout – *Oncorhynchus mykiss* W. 1792) were placed in the other cage. On days two, three, and four of each trial, a minimum of 200 g fish per species (exception: *P. phoxinus* on day one in August (196 g) and in November (183 g)) were placed in the cages, corresponding to two to five individuals per salmonid species and 40 to 90 *P. phoxinus* individuals. At the end of each sampling day, fish were removed from the cages and exchanged for new individuals to generate true biological replicates, prevent acclimation to the local conditions, and reduce stress-periods for the animals. The fish were kept in separate tanks afterwards to avoid the multiple use of individuals. Due to their life cycle and therefore a lack of catchable individuals, the availability of *P. phoxinus* was limited in November and the same individuals had to be used on all sampling days.

Two-liter water samples were used for the detection of fish eDNA. On the first day of each trial (prior to fish placement in cages), control samples were taken from at least five locations between the cage position and the transverse structure. On days two, three, and four, water samples were taken daily at 13 locations downstream of the cages (1.3 km, 555 m, 425 m, 325 m, 225 m, 130 m, 65 m, 33 m, 17 m, 8 m, 4 m, 2.5 m, and 0.5 m; Fig. 1) at approximately the same daytime and before fish were exchanged. Because of changes in the structure of the riverbed, the distance between the cages and the first seven transects sometimes varied by one to two meters between trials, which was accounted for during data analyses. At 0.5 m to 130 m distance, two to four water samples were taken in lateral transects, depending on the width of the river (Fig. 1). Further downstream, we took one sample per location. In November, only two to three lateral transect samples were taken due to limited discharge, resulting in reduced wetted width and sampling at transect 5 had to be omitted due to a slight position change of the cages. Each day, sampling was carried out from the most downstream location moving upstream towards the cages. In September, two sampling runs from 1.3 km to 0.5 m downstream distance were carried out per day.

The water samples were collected right below the stream surface using 2 L wide-neck bottles, which had been treated with chlorine bleach (3.6 g sodium hypochlorite per 100 g liquid) overnight and thoroughly washed using fish-DNA-free tap water. Filtration was carried out on 500 ml filter towers with a peristaltic pump (Solinst; Model 410) and glass fibre filters with 47 mm diameter (1.2 µm mesh width, Whatman GF/C), followed by storage at –80 °C until further processing. In case of filter clogging (high turbidity; only in August), up to three filters were utilized per sample. During all water processing steps, DNA-free gloves were worn and frequently changed. All multi-use equipment was soaked in chlorine bleach for at least ten minutes between samples and thoroughly rinsed using MilliQ water. Forceps for filter handling were singed three times prior to each use. Every ten samples, 2 L of MilliQ water were filtered as negative controls to check for cross-contamination during fieldwork, whereas a sample from the fish transport container served as daily positive control.

Total discharge and its lateral differences were measured with a FlowTracker (Sontek, USA) during the August trial, and with salt and a TQ Tracer (Sommer Messtechnik, Austria) during the November and September trials. In August, discharge increased during the sampling time (10 AM to 5 PM) due to the strong glacial influence but remained constant along the sampling reach. Therefore, diurnal discharge was measured once and supported by point-measurements at 3 PM. In September and November, the inflow of tributaries caused an increase along the 1.3 km sampling reach, with hardly any changes in the course of a day. Hence, measurements were carried out daily directly at the cages, at 225 m, 555 m, and 1.3 km downstream, and supported by a diurnal measurement per trial. Discharge at each water sampling point (in space and time) was interpolated; no rainfall occurred during any of the sampling days. For all three trials, turbidity was measured with a turbidity meter using an infrared light source (AL250T-IR, Aqualytic, Germany). Water temperature, pH, conductivity, and oxygen saturation were obtained with a multi-parameter probe (WTW, Germany).

### Laboratory processing

All molecular work was carried out in a clean-room laboratory at the University of Innsbruck (Austria) compliant with ancient-DNA processing standards. First, filters were lysed using 190 µL TES (0.1 M TRIS, 10mM EDTA, 2% sodium dodecyl sulphate; pH 8) and 10 μL Proteinase K (VWR, 20 mg/mL) each, followed by over-night incubation on a rocking platform at 56°C. Then, filters were transferred to plastic inserts with a perforated bottom and centrifuged for 10 min at 14,000 rpm. The entire resulting lysate was extracted on the Biosprint 96 platform (QIAGEN) using a custom DNA-uptake protocol which combined the DNA contained in up to 900 µL lysate prior to extraction with the “BS 96 tissue protocol” according to the manufacturer’s instructions; except for elution in 100 µL TE. Lysates of filters stemming from the same water sample were combined during the uptake process. Each 96-well plate contained four extraction negative controls and all resulting eluates were stored at -32°C until further processing.

Target DNA was amplified using species-specific primers and celPCR. These were run on Nexus Mastercyclers (Eppendorf); each plate included at least one positive control (DNA extract from target species) and one negative control (molecular grade water). Previously published primers (Thalinger et al., 2016) and a newly developed primer pair for *S. trutta* (Table 1) were utilized. During PCR optimization, primers were tested for specificity against other fish species and aquatic invertebrates occurring in Central European freshwaters (Thalinger et al., 2016). Additionally, we specified assay sensitivity following Sint et al. (2012): reliable positive amplifications were possible for all primer pairs from 10 DNA ds. Each 10 µL PCR master mix contained 1 × Multiplex reaction mix (QIAGEN), 0.5 µM of each primer, 30 mM TMAC, 5 µg BSA (Bovine Serum Albumin) and 3.2 µL DNA extract. Optimized thermocycling conditions were: denaturation at 95°C for 15 min, 35 cycles of 94°C for 30 s, 64°C (66°C for *P. phoxinus*) for 3 min and 72°C for 60 s followed by final elongation at 72°C for 10 min. Target DNA signal strength was determined via capillary electrophoresis on the QIAxcel (QIAGEN) with the associated software QIAxcel Screengel version 1.6.0.10 using the method AM320-30s. The fluorescent signal measured in Relative Fluorescence Units (RFU) was used as a semi-quantitative measure of target DNA (Thalinger, Pütz, & Traugott, 2020; Thalinger et al., 2019) and signals ≥0.08 RFU were deemed positive. To verify the applicability of this approach, 40 samples positive for *S. trutta* were re-tested with digital PCR (Supporting Information (SI) 1a). All extraction and PCR negative controls resulted negative, however, in the August and the November trial six of 16 negative controls showed low levels of contamination (≤0.2 RFU), hence, fluorescence values of potentially affected field samples (10.9% of all obtained PCR results) were down-corrected by the respective values.

**Table 1:**
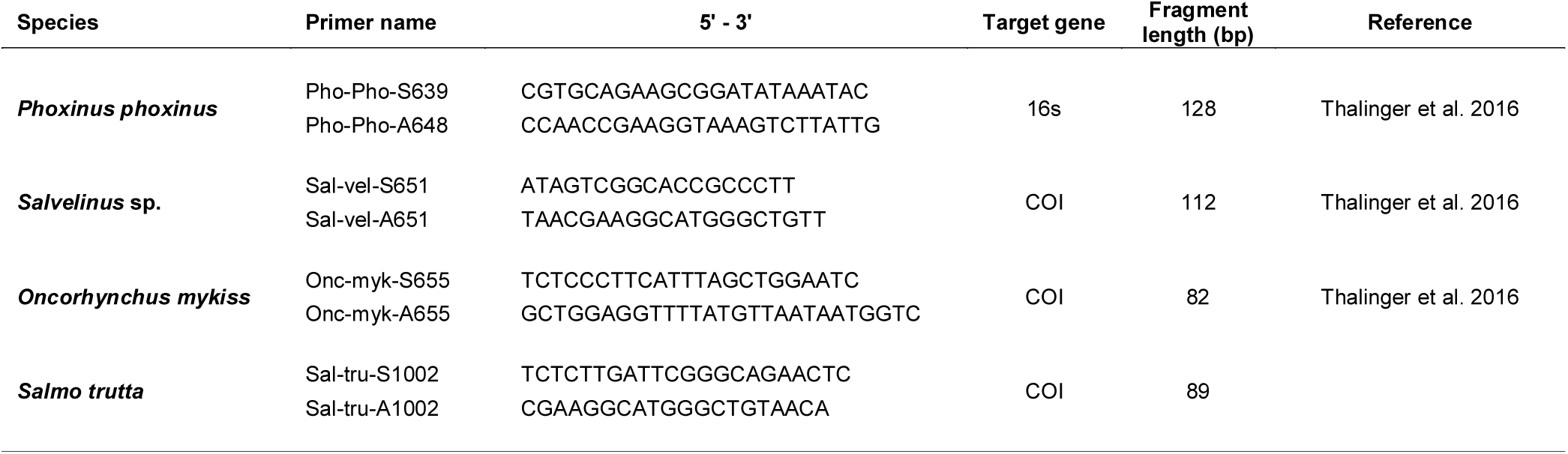
Primer pairs used for the molecular analysis of the eDNA samples. The respective target taxon, primer sequence, target gene, fragment length in base pairs and the source for previously published primers are displayed. Please note that the *Salvelinus* sp. primer pair was designed to amplify both *S. fontinalis* and *S. umbla*.

### Statistical analysis

All data were analyzed in R (R Core Team, 2020) and visualized with “ggplot2” (Wickham, 2016), “ggpubr” (Kassambara, 2019), and “viridis” (Garnier, 2018). We generated species-specific eDNA heatmaps for each trial until 65 m downstream distance and subsequently heatmaps per fish cage, i.e. for salmonids and *P. phoxinus*, for each of the three trials up to 20 m. The lateral distance from the orographically left shore (x), the longitudinal distance from the respective cage (y) and mean RFU (z) were used for the linear interpolation of irregular gridded data with the “akima” package (Akima & Gebhardt, 2016). eDNA signals were not extrapolated towards the edge of the water body and RFU were interpolated on a 5 × 5 cm grid.

Differences in detection rates and eDNA signal strengths amongst salmonid species up to 33 m downstream distance were examined per trial with Z-tests, Kruskal-Wallis tests, and Holm-Sidak-corrected p-values. As no significant differences were detected (SI 1b), salmonid data remained pooled for the subsequent analysis of small-scale lateral and longitudinal eDNA distribution using the packages “reshape2” (Wickham, 2007), “ade4” (Bougeard & Dray, 2018; Chessel, Dufour, & Thioulouse, 2004), and “data.table” (Dowle & Srinivasan, 2020). Lateral changes in interpolated eDNA signals were examined per cage and trial at 2.5 m, 5 m, 10 m and 20 m by testing salmonid and *P. Phoxinus* data for normality and calculating their Spearman Rank Correlation and Holm-Sidak-corrected p-values. To investigate the magnitude of longitudinal eDNA signal changes between 2.5 m and 20 m, differences between eDNA signals of adjacent interpolated datapoints were calculated 1 m from the left shore, at the stream center, and 1m from the right shore individually for each trial, and salmonids and *P. phoxinus*, respectively. Differences were tested for normality and per position, Kruskal-Wallis tests followed by pairwise Wilcoxon-Rank-Sum tests were used to compare between trials for each position (left, center, right) and fish cage (salmonids, *P. phoxinus*); all p-values were Holm-Sidak-corrected.

To analyze the relationship between eDNA signals, distance from the cages, and discharge on a larger scale, only lateral transects with homogenous eDNA distribution at all trials were considered. Therefore, salmonid RFU were tested for normal distribution with a Shapiro-Wilk test, followed by Kruskal-Wallis tests per transect and trial. *P. phoxinus* data were not considered as the dataset was too small for separate testing and mirrored lateral eDNA distributions between the two cages would potentially mask patterns, if considered together. Based on these tests (SI 1c), only data with downstream distances ≥ 130 m were used for the two subsequent analyses: once entering detections as binary variable (0/1), and once using RFU of positive samples and considering non-detections as random results induced by the sampling process (Hagenaars & McCutcheon, 2002). To account for the binary nature and the right-skewed distribution of RFU, Generalized Linear Models (GLM) were used with family “binomial” and a logit-link for detections (0/1) and family “Gamma” with a log-link for RFU (Faraway, 2016). After initial data inspection, discharge, downstream distance, their interaction and fish species were entered as independent variables in both models, and *P. phoxinus* was set as base category (see SI 2 for the complete R code). For both models, simulated, scaled residuals were calculated (package: “DHARMa” (Hartig, 2020); function: “simulateResiduals”; n = 1,000) to confirm model fit and test for outliers and overdispersion.

## Results

Altogether, 306 water samples were analyzed: 84, 168, and 54 from the August, September, and November trial, respectively, and average discharge during sampling 0-20 m downstream of the cages was 835 L/s in August, 174 L/s in September, and 61 L/s in November (maximum 1,593 L/s in August; minimum 59,3 L/s in November). Fifty percent of the 1,224 PCRs resulted positive with the majority of detections occurring in September and November. The environmental variables turbidity (0 – 4.6 NTU), oxygen saturation (9.43 – 11.25 mg/L), water temperature, (4.2 – 10.5°C), conductivity (49.3 – 69.2 µS/cm), and pH (7.11 – 7.6) were near-constant between the three trials. Only in August during sampling within 100 m of the cages (i.e. at a later daytime) turbidity increased (88.7 NTU ± 50.3 NTU SD) and conductivity decreased (36.5 µS/cm ± 9.70 µS/cm SD) because of glacial run-off. Within the first 33 m downstream of the cages, RFU per trial did not differ between the three salmonid species (1 < p > 0.11; SI 1b), leading to a combined analysis of salmonid eDNA signals on a small scale: heatmaps of eDNA signals up to 20 m downstream of the cages revealed eDNA plumes below the respective fish cage and much weaker or no signals at small lateral distances of ∼1 m (Fig. 2; see SI 3 for species-specific heatmaps until 65 m downstream distance). In August, cages were placed in the center of the stream and positive samples within 20 m distance were almost exclusively obtained from sampling points with no lateral offset (Fig. 1 and Fig. 2). The interpolated eDNA signals of the salmonids and *P. phoxinus* obtained in August showed significant positive correlations at all examined distances (2.5 m, 5 m, 10 m, and 20 m; p < 0.001; Table 2). In September, the main water flow out of the salmonid cage was observed towards the orographically right edge of the river; the main water flow from the *P. phoxinus* cage was on the opposite side. This situation was mirrored by high target eDNA signals downstream of the respective cage, low eDNA signals at the opposite sides, and significantly negative correlations between salmonid and *P. phoxinus* RFU at 2.5 m (p < 0.001), 5 m (p < 0.05) and 10 m (p < 0.001; Table 2). In November, eDNA signals were highest for both salmonids and *P. phoxinus* and the eDNA distribution pattern was back-to-front in comparison to September: the main water flow was observed on the orographically left side below the salmonid cage and the orographically right side below the *P. phoxinus* cage leading to significant negative correlations at 5 m (p < 0.001) and 10 m (p < 0.001; Table 2). The small-scale longitudinal changes of the interpolated eDNA signals at the left, center, and right position differed significantly between trials for both salmonids and *P. phoxinus*; exceptions were salmonid signals from the right position and *P. phoxinus* signals from the left position, which did not differ significantly between August and September, respectively (Table 3). The longitudinal changes were strongest in November, with e.g. *P. phoxinus* values decreasing on average 0.036 RFU ± 0.05 RFU SD with each meter; the smallest changes were detected for salmonids in August at the left and right position (Table 3, Fig. 2).

**Figure 2:**
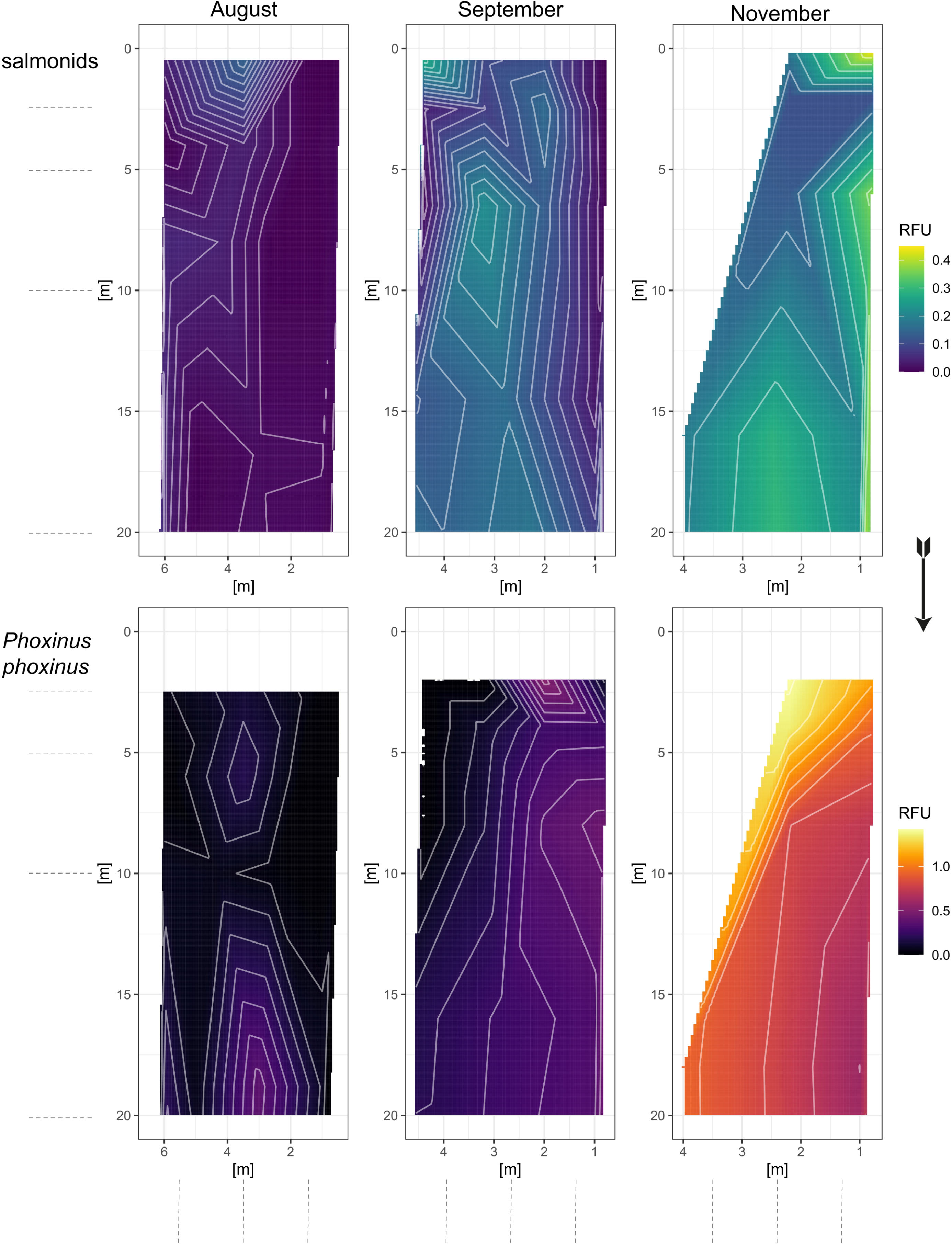
Heatmaps showing the small-scale (≤20 m) distribution of eDNA downstream of the salmonid cage (upper panel) and *Phoxinus phoxinus* cage (lower panel) during August, September, and November. The arrow indicates the flow direction and the cages were placed at zero on the y-axis. As eDNA signals differed between the salmonids and *P. phoxinus*, different colour scales where used between these two groups. Isotherms display interpolated differences of 0.05 RFU. Dashed lines to the left and the bottom of the figures show locations of lateral and longitudinal transects used to compute small-scale differences in eDNA signals; these transects differ from the actual water sampling locations. Note, that the irregular shape in November reflects a gradual increase in wetted width downstream of the cages.

**Table 2:** The correlation between interpolated eDNA signals from the two fish cages. The test statistic (W) and corresponding p-values of Shapiro-Wilk tests for normality are provided per trial, fish cage, and for four downstream distances (2.5 m, 5 m, 10 m and 20 m). Spearman’s rho and the corresponding p-value were calculated per distance and trial. All displayed p-values were Holm-Sidak-corrected.

| trial | longitudinal position | W<br>salmonid<br>cage | p-value | W<br>*P. phoxinus*<br>cage | p-value | Spearman's rho | p-value |
| --- | --- | --- | --- | --- | --- | --- | --- |
| August | 2.5 m | 0.89 | <0.001 | 0.96 | < 0.12 | 0.89 | <0.001 |
|  | 5 m | 0.94 | <0.01 | 0.92 | <0.001 | 0.84 | <0.001 |
|  | 10 m | 0.84 | <0.001 | 0.93 | <0.001 | 0.95 | <0.001 |
|  | 20 m | 0.61 | <0.001 | 0.97 | 0.40 | 0.62 | <0.001 |
| September | 2.5 m | 0.97 | 1.00 | 0.85 | <0.001 | 0.64 | <0.001 |
|  | 5 m | 0.92 | <0.01 | 0.87 | <0.001 | -0.35 | <0.05 |
|  | 10 m | 0.92 | <0.01 | 0.91 | <0.001 | -0.52 | <0.001 |
|  | 20 m | 0.87 | <0.001 | 0.95 | 0.21 | -0.31 | 0.08 |
| November | 2.5 m | 0.42 | <0.001 | 0.96 | 1.00 | 0.30 | 1.00 |
|  | 5 m | 0.84 | <0.01 | 0.91 | 0.19 | -0.87 | <0.001 |
|  | 10 m | 0.82 | <0.001 | 0.76 | <0.001 | -0.82 | <0.001 |
|  | 20 m | 0.96 | 0.67 | 0.95 | 0.27 | -0.26 | 0.49 |

**Table 3:**
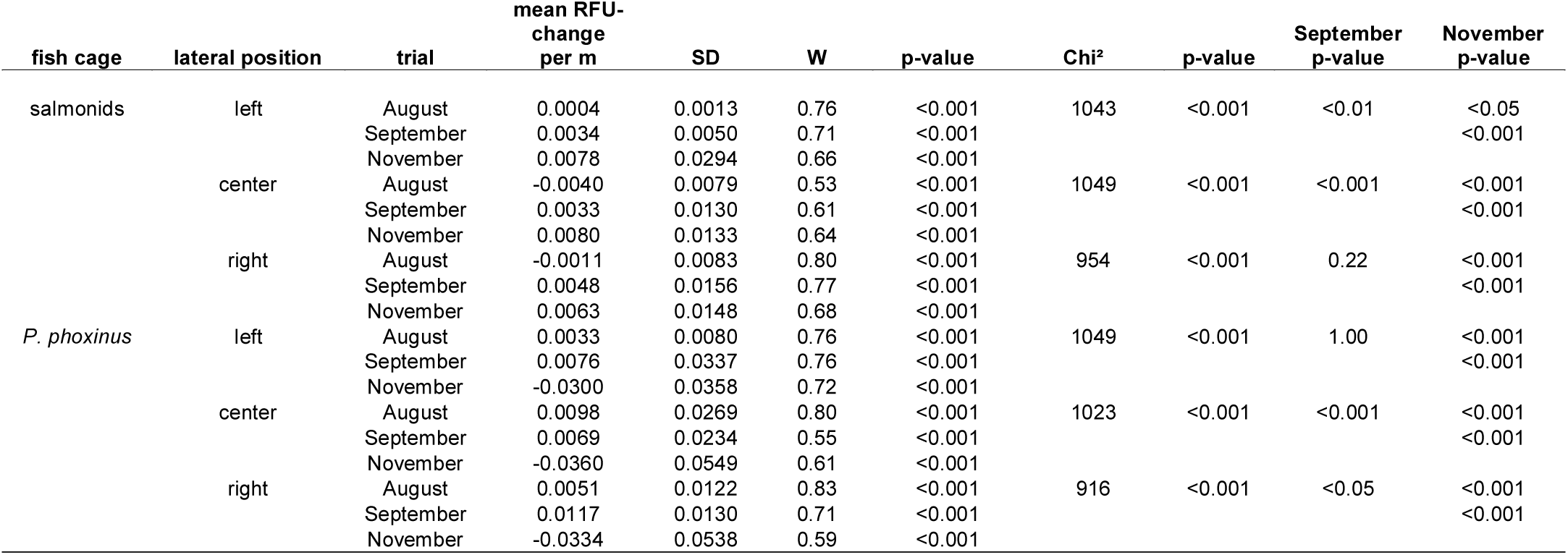
The difference in longitudinal eDNA signal change for the left shore, center, and right shore of the river. The change was compared between the three trials per position and fish cage. The test statistic (W) and p-value of Shapiro-Wilk tests for normality are provided together with the Chi^2^ and p-values obtained from Kruskal-Wallis tests per position and fish-cage. The columns “September p-value” and “November p-value” denote for the results of pairwise Wilcoxon-Rank-Sum tests between individual trials. All displayed p-values were Holm-Sidak-corrected.

At 130 m downstream distance from the cages and beyond, salmonid signals were laterally homogenous during all three trials (SI 1c) and thus used to examine large-scale patterns of detection rates and RFU. In August, only few samples resulted positive with low mean values (e.g. *P. phoxinus* 0.25 RFU ± 0.16 RFU SD; Fig. 3 and Fig. 4). In September, the signals remained at a similar level, but 100% of *P. phoxinus* PCRs and 71% of salmonid species PCRs resulted positive. In November, mean *P. phoxinus* eDNA signals were highest (0.34 RFU ± 0.22 RFU SD), and salmonid signals remained similar to September (Fig. 3 and Fig. 4). The longitudinal increase in discharge was similar in September and November with a 2.1-fold increase from 201 L/s to 335 L/s and a 2.7-fold increase from 71 L/s to 167 L/s, respectively (Fig. 3). Both GLMs (for binomial data and RFU) passed the diagnostics for model fit, outliers, and overdispersion. They displayed similar patterns of parameter estimates and significances (Table 4 and Table 5): salmonids were significantly less likely to be detected and had significantly lower RFU in comparison to *P. phoxinus*. Discharge and downstream distance were significantly correlated in both models and only the individual effect of discharge was not significant (p>0.05) in the Gamma GLM (Table 5).

**Figure 3:**
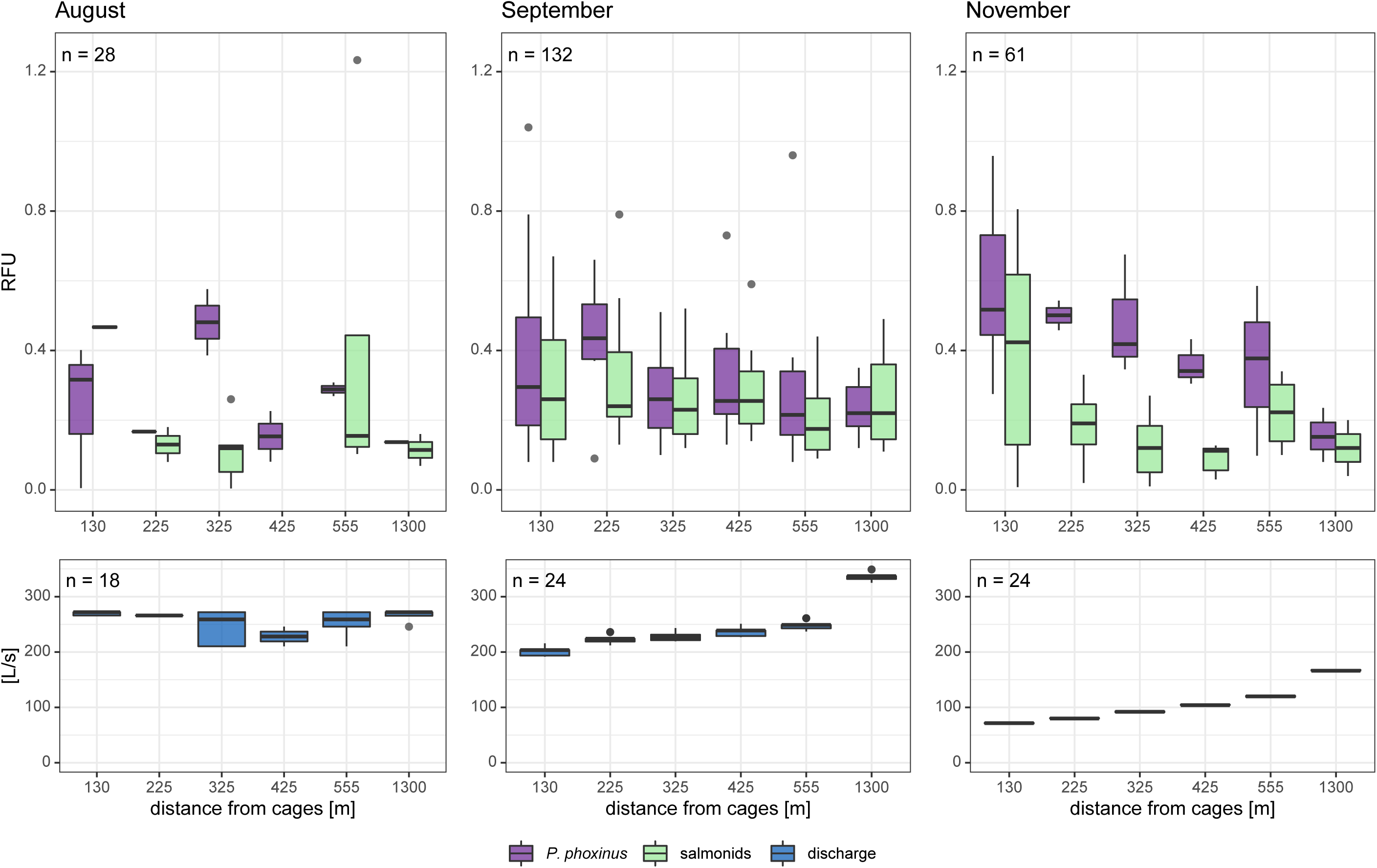
eDNA signal strength of *Phoxinus phoxinus* and salmonids (upper panel) as well as the corresponding discharge (lower panel) determined at different distances from the cages in August, September, and November. The eDNA signal strength was measured in Relative Fluorescence Units (RFU) and is displayed for downstream distances with homogeneous lateral eDNA distribution (≥130 m). Samples testing negative were not considered for this illustration.

**Figure 4:**
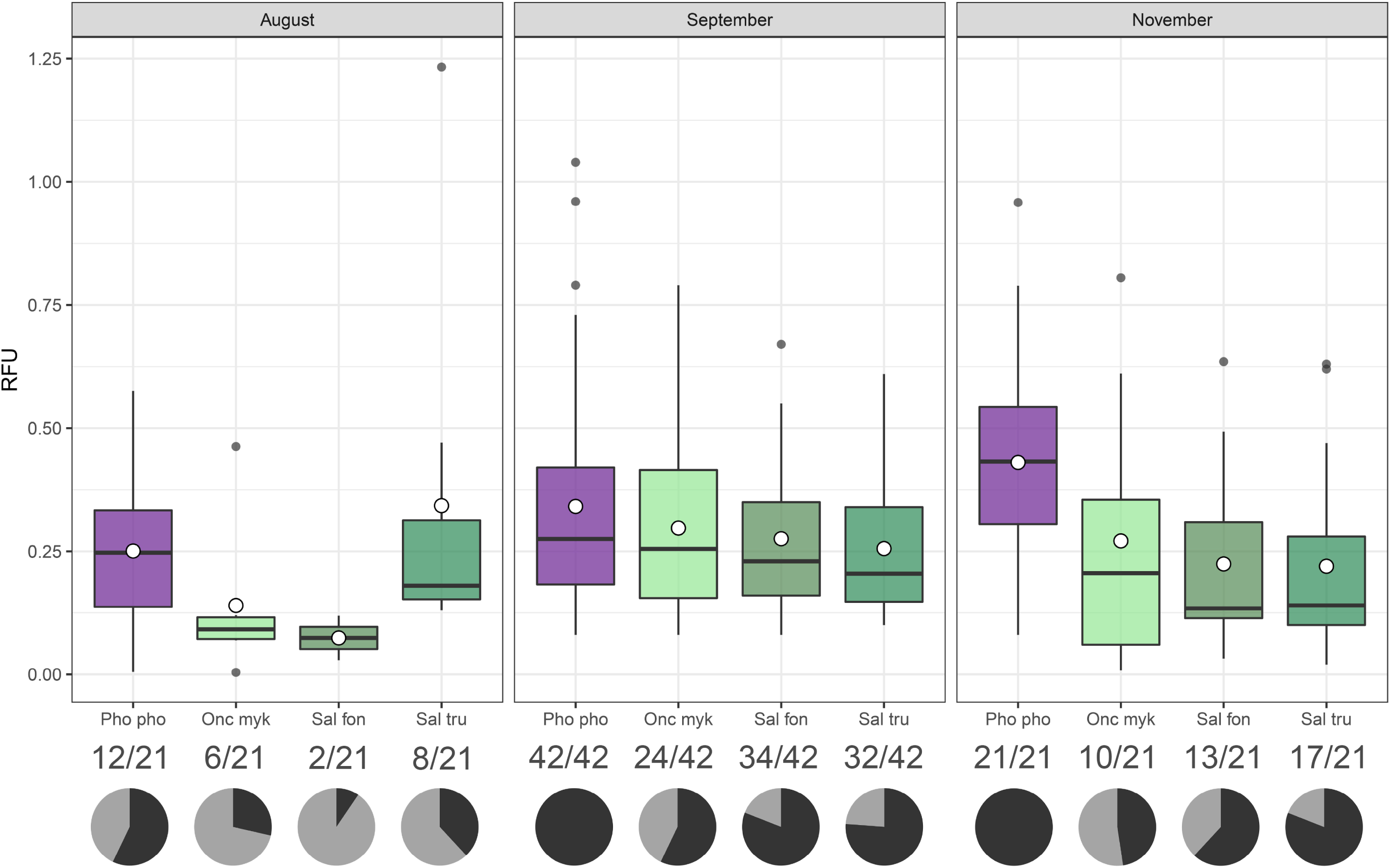
Overall eDNA signal strength measured in Relative Fluorescence Units (RFU) considering water samples collected at distances ≥130 m in August, September, and November. The RFU of positive samples are displayed as boxplots for each fish species individually; the empty circles code for the mean RFU. Underneath each boxplot, the number of samples testing positive / the total sample number are displayed. The pie charts show the proportion of PCR positives for *P. phoxinus* (“Pho pho”) and the three salmonid species (“Onc myk” = *Oncorhynchus mykiss*, “Sal fon” = *Salvelinus fontinalis*, “Sal tru” = *Salmo trutta*) per trial with dark shading coding for successful detection.

**Table 4:** The Generalized Linear Model (Binomial family, logit-link) describing the relation between positive detections and discharge, downstream distance, their interaction, and fish species during the three cage trials. A) The columns show the test statistics (z-value), the lower and upper 95% confidence intervals, and p-values returned from the GLM for the input variables. Table B) shows the degrees of freedom (Df) residuals and deviance for each variable. Note that *Phoxinus phoxinus* was used as base category for the description of species-specific effects.

| A) | Estimate | Std. Error | z-value | lower<br>95% CI | upper<br>95% CI | p-value |
| --- | --- | --- | --- | --- | --- | --- |
| (Intercept) | 4.64 | 0.79 | 5.91 | 3.17 | 6.26 | <0.001 |
| discharge | -0.01 | 0.00 | -3.42 | -0.02 | 0.00 | <0.001 |
| distance | 0.00 | 0.00 | -3.39 | -0.01 | 0.00 | <0.001 |
| *Oncorhynchus mykiss* | -2.32 | 0.43 | -5.45 | -3.21 | -1.52 | <0.001 |
| *Salvelinus fontinalis* | -1.86 | 0.43 | -4.37 | -2.75 | -1.06 | <0.001 |
| *Salmo trutta* | -1.43 | 0.43 | -3.31 | -2.32 | -0.61 | <0.001 |
| discharge*distance | 0.00 | 0.00 | 3.32 | 0.00 | 0.00 | <0.001 |
| B) | Df | Deviance | Resid. Df | Resid. Dev |  |  |
| NULL |  |  | 335 | 431.78 |  |  |
| discharge | 1 | 3.728 | 334 | 428.05 |  |  |
| distance | 1 | 0.433 | 333 | 427.62 |  |  |
| fish species | 3 | 39.212 | 330 | 388.4 |  |  |
| discharge*distance | 1 | 12.017 | 329 | 376.39 |  |  |

**Table 5:** The Generalized Linear Model (Gamma-family, log-link) describing the relation between RFU and discharge, downstream distance, their interaction, and fish species during the three cage trials. A) The columns show the test statistics (t-value), the lower and upper 95% confidence intervals, and p-values returned from the GLM for the input variables. Table B) shows the degrees of freedom (Df) residuals and deviance for each variable. Note that *Phoxinus phoxinus* was used as base category for the description of species-specific effects.

| A) | Estimate | Std. Error | t-value | lower<br>95% CI | upper<br>95% CI | p-value |
| --- | --- | --- | --- | --- | --- | --- |
| (Intercept) | -0.42 | 0.22 | -1.96 | -0.82 | -0.01 | 0.05 |
| discharge | 0.00 | 0.00 | -1.74 | 0.00 | 0.00 | 0.08 |
| distance | 0.00 | 0.00 | -4.21 | 0.00 | 0.00 | <0.001 |
| *Oncorhynchus mykiss* | -0.33 | 0.13 | -2.52 | -0.58 | -0.07 | <0.05 |
| *Salvelinus fontinalis* | -0.38 | 0.12 | -3.14 | -0.62 | -0.14 | <0.01 |
| *Salmo trutta* | -0.30 | 0.12 | -2.61 | -0.53 | -0.07 | <0.01 |
| discharge*distance | 0.00 | 0.00 | 3.25 | 0.00 | 0.00 | <0.01 |

| B) | Df | Deviance | Resid. Df | Resid. Dev |
| --- | --- | --- | --- | --- |
| NULL |  |  | 220 | 125.14 |
| discharge | 1 | 1.4257 | 219 | 123.72 |
| distance | 1 | 3.8624 | 218 | 119.85 |
| fish species | 3 | 4.9894 | 215 | 114.86 |
| discharge*distance | 1 | 5.0312 | 214 | 109.83 |

## Discussion

Our results demonstrate profound spatio-temporal changes in eDNA distribution induced by seasonal discharge conditions. Until 20 m the lateral position of the eDNA plumes downstream of the caged fish mirrored their position inside the river with salmonid and *P. phoxinus* plumes in the stream center in August and on opposing sides in September and November. Small-scale longitudinal changes in eDNA signals were strongest at low discharge in November, and lateral mixing occurred further downstream at higher discharge in August when significant lateral differences were detected at 65 m. Additionally, the small-sized *P. phoxinus* individuals emitted a significantly stronger eDNA signal and had higher detection rates than the three larger-bodied salmonid species, principally in November. Along the 1.3 km flow path, increasing discharge and distance both had significantly negative effects on detection rates and eDNA signal strength.

The eDNA signals recorded at a small lateral and longitudinal scale (≤20 m) coincide with previous results describing a plume-shaped distribution of eDNA downstream of the source (Laporte et al., 2020; Wilcox, McKelvey, Young, Lowe, & Schwartz, 2015; Wood et al., 2020). Additionally, the shape of the salmonid and *P. phoxinus* plume (averaged across three sampling days) mirrored the lateral position of the respective cage when these were not aligned in flow direction in September and November. This finding complicates the interpretation of field-derived eDNA results when local conditions differ across the wetted width, especially as individual species may prefer specific micro-habitats depending on hydrological conditions (e.g. seek shelter from high flow velocities) (Aarts & Nienhuis, 2003). Based on our experimental setup with constant fish biomasses, it was not possible to confirm the results of Wood et al. (2020) who found a positive correlation between eDNA quantities and detection probability at higher lateral distance. However, our data show a comparable effect induced by lower discharge and slower flow velocities (Wondzell, Gooseff, & McGlynn, 2007) at constant fish biomass. Future studies examining lateral and longitudinal eDNA distribution should therefore take biomass-induced variations and hydrological effects into account.

All three salmonid species (average individual mass 101 g) had lower detection probabilities and lower eDNA signals than *P. phoxinus* (average individual mass 3.5 g) at similar total biomass, a finding which was primarily driven by the results of the trial in November. eDNA shedding rates and biomass in most cases exhibit a positive relationship in controlled and natural settings (Lacoursière-Roussel, Rosabal, & Bernatchez, 2016; Takahara, Minamoto, Yamanaka, Doi, & Kawabata, 2012; Thalinger et al., 2019). Maruyama et al. (2014) detected elevated levels of eDNA release per gram bodyweight in juvenile fish and recently, allometrically scaled fish mass was found to best describe the relationship between eDNA signals and brook trout populations in lakes (Yates et al., 2020). In the present study, several scenarios can explain the difference in detection probability and signal strength: the *P. phoxinus* individuals, although mostly not juvenile, could indeed have had higher metabolic rates (Vinberg, 1960), which are common for smaller fish species (Clarke & Johnston, 1999). The increased surface to volume ratio of the smaller fish is another potential explanation as more *P. phoxinus* surface was exposed to the water flow. The eDNA release rates could generally differ between cyprinids and salmonids, which also exhibit structural and physiological differences (Freyhof & Kottelat, 2007). Albeit the daily exchange of fish individuals after eDNA sampling was another potential source of variation (e.g. slight changes in total biomass or individual counts) we deemed this measure appropriate to obtain true biological replicates, avoid acclimation effects to the stream environment, and minimize the exposure to stressful flow conditions. Generally, the salmonids were better adapted to the high flow conditions in August, when some *P. phoxinus* did not survive the exposure to 1,500 L/s discharge. This could explain the distinct detection probability for *P. phoxinus*, as dead individuals can emit more eDNA into the water (Tillotson et al., 2018). However, the difference in eDNA signal strength was principally found in November at ∼60 L/s, when all but one individual of *P. phoxinus* survived. Though celPCR assay optimization (Sint et al., 2012; Thalinger et al., 2016) can not fully account for differences in amplification efficiency between primer pairs (Thalinger et al., 2020), primer bias is an unlikely cause for the strong *P. phoxinus* signals, because detection patterns and signal strength did not differ between the salmonids. Nevertheless, celPCR produces a semi-quantitative measure of target copy numbers and without direct comparison to a fully quantitative approach (i.e. digital PCR), it is not possible to infer absolute eDNA concentrations from RFU (Thalinger et al., 2020). Most likely, the strong eDNA signals of *P. phoxinus* can be attributed to a combination of the aforementioned physiological and morphological differences. These effects could be even stronger when fish are compared to other aquatic groups such as amphibians, mussels, or crayfish (Bedwell & Goldberg, 2020; Robinson, Uren Webster, Cable, James, & Consuegra, 2018; Wacker et al., 2019).

As expected for glacier-fed Alpine rivers and streams, discharge changed substantially (25-fold) between trials (Bard et al., 2015). The associated diluting effect was clearly visible from eDNA signal strengths directly downstream of the cages (≤20 m): average *P. phoxinus* signals at this location ranged from 0.30 RFU ± 0.30 RFU SD in August to 1.06 RFU ± 0.46 RFU SD in November. Few of the samples taken at high discharge >1,000 L/s in August tested positive, but in this situation the 200 g fish mass per species were less than 0.2 ‰ of the passing water mass. The summer situation was most challenging to examine, as discharge and turbidity increased from morning (∼270 L/s and clear water) to afternoon (∼1,100 L/s and high turbidity). The analysis of samples with high variations in turbidity should include this variable in the modelling process to account for changes in extraction efficiency and inhibition associated with increased turbidity (Harper, Buxton, et al., 2019). We refrained from doing so as the samples taken in August at high turbidity were not part of the dataset used for modelling. In contrast to classic fish monitoring via electrofishing, eDNA-based methods are not restricted to low flow conditions outside of protected periods and spawning seasons. Based on our results, spring and summer sampling in glacier-fed Alpine rivers should, however, not take place at high discharge in the afternoon and evening and is also not advisable during floods and after strong rainfalls especially without increasing the sample number or the filtered water volume (Laporte et al., 2020). In September and November discharge remained almost constant over time but increased more than two-fold (tributary inflow) within the examined 1.3 km. So far, most eDNA studies only discuss the effect of changing discharge (Harrison et al., 2019; Laramie, Pilliod, & Goldberg, 2015; Wood et al., 2020) even though it generally increases along the course of running waters and significantly influenced eDNA detection rates this study. Furthermore, the significance of the discharge-distance interaction term in our analysis shows the effect of the distinct hydrological conditions at each trial on eDNA detection rates and signal strengths.

DNA signals and detection rates of both *P. phoxinus* and the salmonids declined significantly with increasing distance from their source, as expected from previous work on eDNA deposition and degradation (Harrison et al., 2019). At constant longitudinal and temporal discharge, average transport distance (*SP*) and depositional velocity (*vdep*) are used to describe eDNA deposition (Pont et al., 2018; Shogren et al., 2017; Wilcox et al., 2016). The calculation of *vdep* relies on flow velocity and depth data, which can fluctuate considerably in natural and semi-natural rivers and are directly influenced by discharge. Therefore, we refrained from calculating this factor. *SP* is commonly described as the slope parameter of a first order exponential decline *SP = 1/-k* (but also see Wood et al., 2020). In our case this would result in an average transport distance of ∼ 3,000 m and ∼850 m for *P. phoxinus* eDNA in September and November, respectively (based on data from locations with homogenous lateral eDNA distribution), and confirm the positive correlation between river size in general and transport distance (Deiner & Altermatt, 2014; Jane et al., 2015; Pont et al., 2018; Shogren et al., 2017). However, *SP* is non-generalizable between streams (Harrison et al., 2019) and thus of limited use outside of a systematic framework incorporating the complex flow regime of Alpine rivers.

Our work aids to the understanding of how eDNA signals obtained from field-collected samples can be interpreted for species monitoring and conservation. Sampling campaigns carried out in dynamic habitats such as Alpine rivers and streams need to account for the heterogenous lateral eDNA distribution, adapt the sampling scheme to habitat preferences of the target species, and address the prevailing discharge situation. In a best-case scenario, the target species has distinct habitat preferences, and discharge is low and constant during the entire sampling period. Then, eDNA quantities measured directly downstream of suitable habitats are likely to be directly correlated with local target species biomass (Hinlo, Lintermans, Gleeson, Broadhurst, & Furlan, 2018). Otherwise, only discharge measurements at each sampling location can prevent flawed inferences (Thalinger et al., 2019). The detection of rare species, however, is best accomplished by determining a suitable distance between sampling points and preliminary investigations of the respective eDNA shedding rates (cp. Wood et al., 2020). The population densities and the position of individuals within lotic systems should not be inferred from eDNA signals without any *a priori* knowledge on local hydrology. Comparisons between species are also not advisable without previous tests under controlled conditions. Therefore, we advocate for the reporting of the fluvial morphology (e.g. pool, riffle, run), local discharge, time, and species biology during field sampling. In the future, these data can be incorporated in hydrological models specifically designed for eDNA-based species monitoring.

## Data Availability Statement

All data on fish, discharge, eDNA signals, environmental conditions, sampling and comparison between celPCR and dPCR have been uploaded to Figshare and are available at https://doi.org/10.6084/m9.figshare.12380642.v1

### Acknowledgements

This research was conducted within the eDNA-Alpfish project funded by the Austria Research Promotion Agency (FFG); project number 853219. We thank F. Drewes for his extraordinary patience, support and organisational skills during setting up the cages, fish handling, and discharge measurements in the face of non-existent mobile reception. We are grateful to H. Raffl for letting us conduct the experiments in his fishing area and M. Böcker for supporting this work with her graphic skills. We thank two anonymous reviewers for their constructive feedback on the original manuscript.

## Conflict of interest

MT is the co-founder of Sinsoma GmbH, a for-profit company dedicated to DNA analyses in environmental studies; CM and RS are co-founders of the ARGE Limnologie GesmbH, a for-profit consultancy in aquatic ecology.

## Author contributions

MT, JW, CM, and RS conceived the study; the experiment was designed by BT, MT, RS, and CM. Data were acquired and analyzed by BT, DK, and YP. BT wrote the first draft of the manuscript which was revised by DK, YP, JW, and MT.

## Notes

https://doi.org/10.6084/m9.figshare.12380642.v1

