## Supplementary material for "Lateral and longitudinal fish eDNA distribution in dynamic riverine habitats": SI 1

**Supporting Information 1**

**Supporting Information 1a: Comparison of digital PCR and celPCR**

To confirm the suitability of endpoint PCR coupled with capillary electrophoresis (celPCR) for semi-quantitative estimations of target DNA, 40 samples from the caged fish experiment which tested positive for *Salmo trutta* were re-run on a digital PCR system (dPCR; Bio-Rad). Each reaction contained 1 × EvaGreen Supermix (Bio-Rad), 220 nM of *S. trutta* forward and reverse primer, 5.28 µl DNA extract and PCR-grade water to obtain the total volume of 22 µl of which 20 µl were used for droplet generation on the AutoDG (Bio-Rad) with Droplet Generation Oil for EvaGreen (Bio-Rad). Optimized dPCR conditions were: denaturation at 95 °C for 5 min, followed by 40 cycles of 95 °C for 30 s, 62 °C for 1 min and 72 °C for 1 min, and final stabilization at 4 °C for 5 min followed by 90 °C for 5 min. Target fluorescence was measured on the QX200 Droplet Reader (Bio-Rad) using the Quanta Soft Analysis Pro Software 1.0.596 (Bio-Rad). All dPCR results are based on more than 13,000 droplets per reaction. The obtained target DNA concentrations were correlated with the respective Relative Fluorescence Units (RFU) obtained from celPCR and showed a linear relationship (SI 1 Fig. 1).

All data are available at Figshare (<https://doi.org/10.6084/m9.figshare.12380642.v1>)

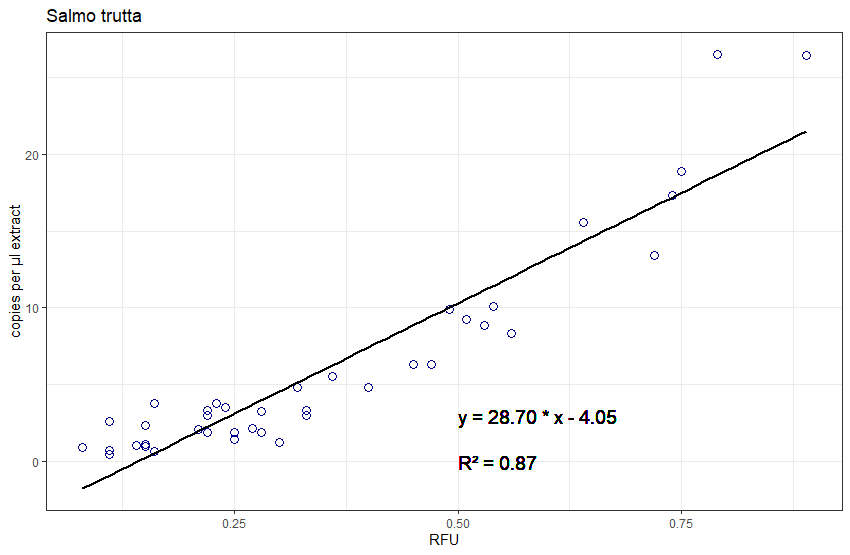

**SI 1 Fig. 1:** The linear relationship between target copies measured with dPCR and (RFU) measured with celPCR. Data points, regression line, and equation are displayed for 40 field samples from the caged fish experiment. P-values of the slope and intercept parameter are < 0.001.

**Supporting Information 1b: Differences between salmonids <33 m downstream distance**

Differences in detection rates and eDNA signal strengths among salmonid species up to 33 m downstream distance were examined for each of the three trials with Z-tests, Kruskal-Wallis tests, and Holm-Sidak-corrected p-values (SI 1 Table 1).

**SI 1 Table 1:** eDNA signals obtained from the three salmonid species tested for differences in detection probability and RFUs.

| **trial** | **X²** | **p-value** | **Chi²** | **p-value** |
| --- | --- | --- | --- | --- |
| August | 5.84 | 0.11 | 3.71 | 0.47 |
| September | 0.69 | 1 | 1.36 | 1 |
| November | 2.15 | 0.68 | 4.31 | 0.35 |

Per trial, salmonid eDNA signals obtained from the cages until 33 m downstream distance were tested for differences in detection probability using z-tests (test statistic X²); Kruskal-Wallis tests (test statistic Chi²) were used to test for differences in obtained RFUs.

**Supporting Information 1c: Tests for laterally homogenous eDNA distribution**

To analyze the relationship between eDNA signals, distance from the cages, and discharge on a larger scale, only transects with homogenous eDNA distribution at all trials were considered. Therefore, salmonid RFU were tested for normal distribution with a Shapiro-Wilk test, followed by Kruskal-Wallis tests per transect and trial. Based on these tests (SI 1 Table 2), only data with downstream distances ≥ 130 m were used for the two subsequent analyses.

**SI 1 Table 2:** Results of Shapiro-Wilk and Kruskal-Wallis tests with salmonid eDNA signals per transect and trial

| **trial** | **transect** | **W** | **p-value** | **Chi²** | **p-value** |
| --- | --- | --- | --- | --- | --- |
| August | 1 | 0.37 | <0.001 | 6.27 | 0.10 |
|  | 2 | 0.38 | <0.001 | 6.91 | 0.07 |
|  | 3 | 0.43 | <0.001 | 5.96 | 0.11 |
|  | 4 | 0.30 | <0.001 | 1.04 | 0.59 |
|  | 5 | 0.38 | <0.001 | 2.01 | 0.37 |
|  | 6 | 0.50 | <0.001 | 11.66 | <0.01 |
|  | 7 | 0.37 | <0.001 | 0.007 | 0.94 |
| September | 1 | 0.45 | <0.001 | 12.65 | <0.01 |
|  | 2 | 0.59 | <0.001 | 20.98 | <0.001 |
|  | 3 | 0.69 | <0.001 | 37.00 | <0.001 |
|  | 4 | 0.67 | <0.001 | 6.18 | <0.05 |
|  | 5 | 0.80 | <0.001 | 0.09 | 0.96 |
|  | 6 | 0.84 | <0.001 | 0.13 | 0.94 |
|  | 7 | 0.90 | <0.001 | 0.10 | 0.75 |
| November | 1 | 0.93 | 0.21 | 5.33 | <0.05 |
|  | 2 | 0.74 | <0.001 | 0.0002 | 0.96 |
|  | 3 | 0.82 | <0.01 | 3.92 | <0.05 |
|  | 4 | 0.90 | <0.05 | 1.15 | 0.56 |
|  | 6 | 0.95 | 0.44 | 0.007 | 0.93 |
|  | 7 | 0.86 | <0.05 | 0.03 | 0.86 |

Per trial and transect (1 to 7) Shaipro-Wilk test statistic (W) and p-values as well as the corresponding Chi² and p-values obtained from Kruskal-Wallis tests are reported. Please note, that in November transect 5 did not exist.
