## Supplementary material for "Lateral and longitudinal fish eDNA distribution in dynamic riverine habitats": SI 3

**Supporting Information 3**

Heatmaps of small-scale eDNA distribution during each of the three trials (August, September, November) were generated individually for each of the four used fish species: *Phoxinus phoxinus*, *Oncorhynchus mykiss*, *Salmo trutta*, and *Salvelinus fontinalis*. The data were analyzed in R (R Core Team, 2020) using the “akima” package (Akima & Gebhardt, 2016) for the linear interpolation of irregular gridded data. eDNA signals were not extrapolated towards the edge of the water body and interpolated between sampling locations on a 5 × 5 cm grid. The following figures present eDNA distribution downstream of the cages up to a distance of 65 m. Please note that the colour scheme is uniform per trial, but not between trials.

All data are available at Figshare (<https://doi.org/10.6084/m9.figshare.12380642.v1>)

**August**

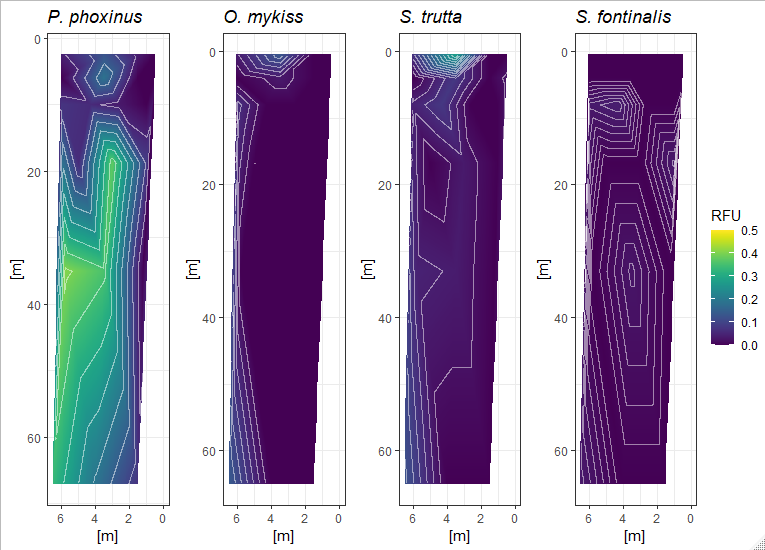

**September**

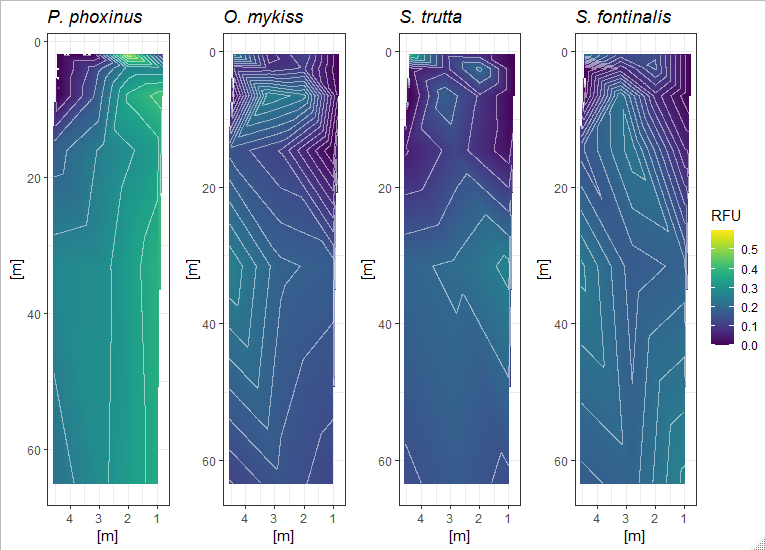

**November**

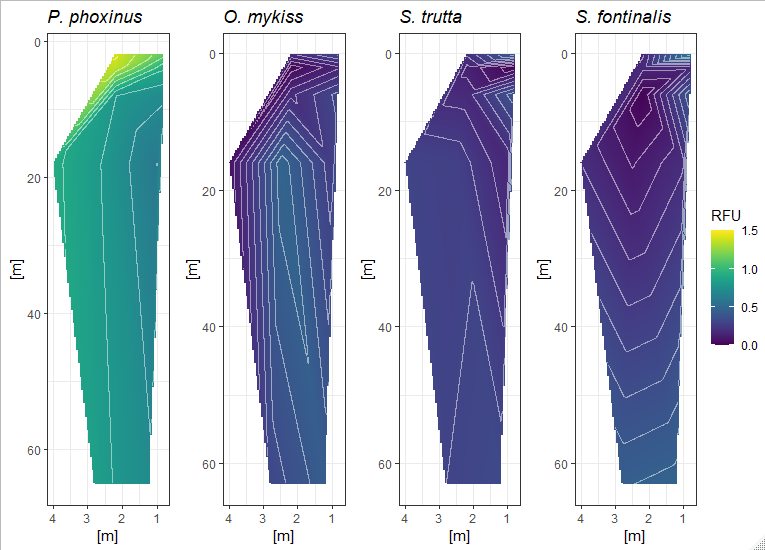
